# Superoxide dismutases shape manganese stoichiometry in Southern Ocean diatoms

**DOI:** 10.1101/2025.09.24.678368

**Authors:** J. Scott P. McCain, Loay J. Jabre, Elden Rowland, Jinyoung Jung, Youngju Lee, Tae-Wan Kim, Corina P. D. Brussaard, Rob Middag, Erin M. Bertrand

## Abstract

Elemental stoichiometry of biomass is a focal point that connects different biogeochemical cycles. Yet, the mechanistic underpinnings of elemental stoichiometry are poorly quantified in many cases. We combined targeted and untargeted metaproteomics, Bayesian statistical modelling, and geochemical measurements to quantify the contribution of specific proteins to metal stoichiometry in natural populations of Southern Ocean diatoms. Our analyses indicate that a substantial amount of non-photosynthetic manganese (Mn) in diatoms in an Antarctic polynya can be attributed to superoxide dismutases (∼0.7 µmol Mn: mol Carbon; ∼20% of the total cellular Mn quota). We then used cultures and proteomic profiling of the key polar diatom *Fragilariopsis cylindrus* to identify environmental controls on superoxide dismutases, and discovered that iron concentration has little influence on the abundance of two Mn superoxide dismutases, while Mn limitation induces the depletion of these Mn superoxide dismutases and an accompanying increase of nickel superoxide dismutase. Overall, we combined metaproteomic approaches to quantify proteomic composition and connected these measurements to metal-to-carbon ratios and their responses to metal availability. Because metal quotas are key parameters in some biogeochemical models, our approach provides a direct mechanism for informing ecosystem-scale models with molecular measurements.

## Main

Elemental stoichiometry is an emergent property of organisms, which plays critical roles in connecting different biogeochemical cycles and influencing dissolved concentrations of elements. A fundamental challenge in biogeochemistry is deconstructing this emergent property. For example, measurements of macromolecular composition (e.g. protein, RNA, DNA, lipid and carbohydrate content) through gradients in nitrogen starvation have begun to identify mechanistic underpinnings of macronutrient elemental ratios^1^. Using these mechanistic insights, global biogeochemical models with explicit representations of macromolecular composition have shown that elemental stoichiometry is driven by various macromolecules^2^.

The metal-to-carbon ratio is an important stoichiometric property of organisms, particularly because trace metals limit primary productivity in large swathes of the ocean. For example, Tagliabue *et al* (2018)^3^ showed using a biogeochemical model that the iron-to-carbon (Fe:C) ratio in phytoplankton might have dramatic impacts on total biomass of upper trophic levels in the eastern Equatorial Pacific. Iron (Fe) and manganese (Mn) are two trace metals that limit primary productivity in the Southern Ocean^4–8^, and most of the focus on deconstructing Fe:C and Mn:C has been on photosynthesis^9,10^, a major sink for these metals. However, there are many other biomolecules which require Fe and Mn (e.g., superoxide dismutases), but we have limited quantitative assessments of how these biomolecules influence metal-to-carbon ratios^11,12^. A major challenge in answering this question is to make precise, molecular measurements under relevant environmental conditions, and then subsequently convert those measurements into biogeochemically relevant quantities like elemental stoichiometry. Successfully making this connection, from molecular to biogeochemical quantities of interest (e.g., elemental quotas or uptake rates), would provide a road to mechanistically justifying parameters in large-scale biogeochemical models (similar to other approaches successfully achieved with macromolecular observations^2^). This connection would provide a new generation of constraints for biogeochemical models leveraging molecular data sources from the global ocean, which have been mostly qualitatively informing biogeochemical models. Overall, biogeochemical models are our best conceptualizations of the dynamic relationship between phytoplankton and the global ocean, and so mechanistically grounding parameterizations is crucial.

The increasing evidence that Mn influences primary production in the Southern Ocean^4–8,13^ warrants mechanistic models that represent how Mn is used by phytoplankton. Previous work has shown that altering parameters and model structures that influence the breakdown of the Mn quota, via the Mn:C biomass, are predicted to have impacts on carbon export^14,15^. Various studies have focused on three main Mn pools: Mn from photosystem II proteins (PSII), Mn from Mn superoxide dismutase (MnSOD), and Mn storage^9,15,16^. One study from a non-polar and non-marine diatom, *Thalassiosira pseudonana*, suggests that MnSOD contributes ∼10-30% of total cellular Mn, while Mn from PSII contributes 50-80%^12^. Given that Mn limitation has only been observed in the Southern Ocean (to our knowledge, to date), there are several key unknowns: What is the contribution of MnSOD to Mn quotas in Southern Ocean diatoms under natural conditions? And, what are the environmental drivers for MnSOD abundances in Southern Ocean diatoms?

Here we combine targeted and untargeted metaproteomic measurements of samples from the Southern Ocean, and Bayesian statistical modelling to empirically describe the contribution of MnSOD to Mn quotas. We specifically focus on quantifying proteomic composition in the numerically abundant Southern Ocean diatom species *Fragilariopsis kerguelensis*. We provide evidence that MnSOD is an important component of the non-photosynthesis-associated Mn quota in natural populations of *Fragilariopsis kerguelensis*. We then show with cultures and proteomic profiling of the related polar diatom *Fragilariopsis cylindrus* that MnSOD abundance is independent of Fe concentrations, which has important implications for mechanistically modelling Mn quotas.

### Combining metaproteomic approaches to quantify taxon-normalized proteomic composition

We collected samples for quantifying proteomic composition of phytoplankton in the Amundsen Sea (Southern Ocean) during the austral summer in 2018 aboard the RV Araon. Water samples from 15 stations (total of 42 samples, ranging from 6 to 68 m depth) were filtered using size-fractionation (0.2, 3.0, and 12.0 µm pore size of a polycarbonate filter) for metaproteomics (Fig. 1A, B). Water was also collected at similar depths using a trace-metal clean CTD^17^ for concentrations of dissolved trace metals (geochemical data have been previously published^18,19^; see Supplementary Fig. S1-S4 for depth profiles of dissolved Mn and Fe, photosynthetically active radiation, and temperature).

**Figure 1.**
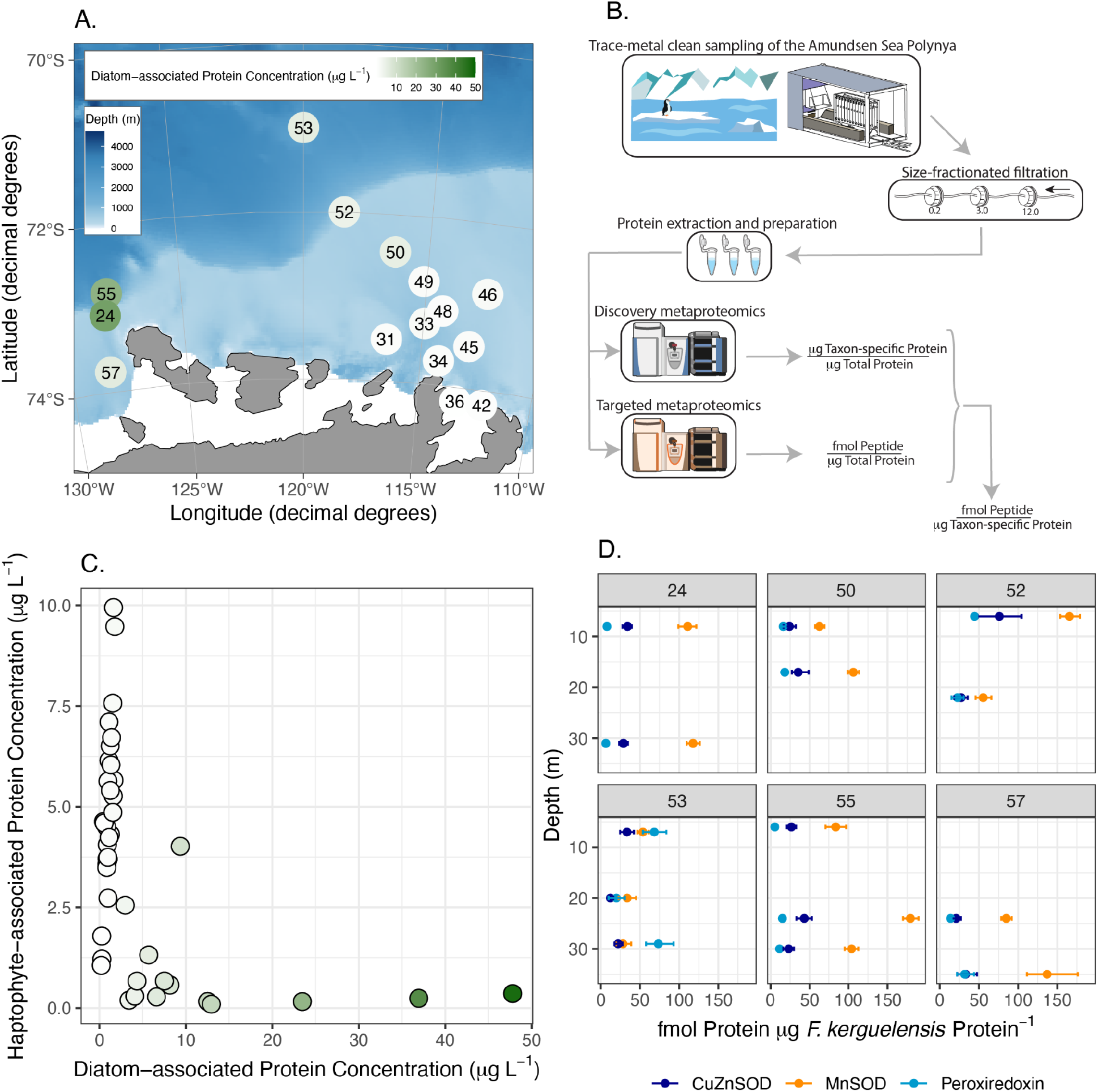
Using metaproteomic sampling of the Amundsen Sea to explore biogeography and proteome composition. (A) Fifteen stations in the Amundsen Sea Polynya and marginal sea ice zone were sampled. Numbers in circles are station numbers, and the colour of the circle represents the depth-averaged diatom-associated protein concentration (scale bar inset). (B) Overview schematic. We sampled water in the Amundsen Sea using trace-metal clean methods. Water was size-fractionated via serial filtration, biomass filters stored frozen and, then protein was extracted and prepared for mass spectrometry. For each 3.0 µm filter we conducted untargeted and targeted mass spectrometry, which allowed us to estimate the fmol of a given peptide per µg protein for our taxon of interest, *Fragilariopsis kerguelensis*. (C) Metaproteome-derived concentrations of haptophytes and diatoms showed geographic separation of these two dominant taxonomic groups. Colours reflect the diatom-associated protein concentration as in (A). (D) Profiles of estimated taxon-normalized protein abundances, CuZnSOD, MnSOD, and peroxiredoxin across six stations where *F. kerguelensis* was abundant. Panel labels refer to the station number.

For each 3.0 µm filter we conducted untargeted and targeted mass spectrometry-based metaproteomics (Materials and Methods), which allowed us to estimate the fmol of a given peptide per μg protein for our taxon of interest, *Fragilariopsis kerguelensis*. Throughout we refer to untargeted metaproteomics as a mass spectrometry method for attempting to identify and quantity all peptides in a given sample (Materials and Methods), and we refer to targeted metaproteomics as a mass spectrometry method for relying on isotopically labeled synthetic peptide standards that are spiked into a sample to quantify endogenous peptide abundances. Untargeted metaproteomics was conducted on the 3.0 µm filter sizes to focus on the diatom populations. We used a previously published metatranscriptome^20^, supplemented with a small set of antioxidant proteins from metagenome-assembled genomes^21^, as a database to search our mass spectra against. We used database-independent normalization, which allowed us to make quantitative statements about the proportion of protein in a sample attributed to a taxonomic group (Methods and Materials).

Combining untargeted metaproteomic measurements and absolute concentrations of protein per litre of seawater provided a basic biogeography of the Amundsen Sea polynya (Fig. 1C; Materials and Methods). Consistent with other observations of phytoplankton in Antarctic polynyas^18,22^, we observed a clear biogeographic separation in this size fraction (between 3 and 12 micron filter pore size), where the open water of the polynya was dominated by the haptophyte *Phaeocystis antarctica* and the marginal sea ice zone (specifically stations 50, 52, 53, 57, 55, and 24; Fig. 1A) by diatoms from the genus *Fragilariopsis*. This pattern is suggestive of niche partitioning via resource competition (e.g., light or nutrients; similar to those observed in freshwater ecosystems^23^). Overall, our results demonstrate the value of untargeted metaproteomics for providing a quantitative biogeographic census at various levels of taxonomic resolution.

*Fragilariopsis kerguelensis* unduly contributes to biomass in some of these samples (Supplementary Fig. S5) and in other regions of the Southern Ocean^24^. This dominance makes *F. kerguelensis* an excellent target system for our central goal to estimate proteomic composition and ultimately connect to elemental stoichiometry. Untargeted metaproteomics can be used to estimate the proportion of protein attributed to a specific taxonomic group^25,26^ (as described above and in Methods; given there are sufficient taxonomically-informative peptides). One unsolved challenge is that there are often small numbers of detected peptides that are both taxonomically- and functionally-informative, fewer than are required to make statements about proteomic composition for a specific species. The reliance on few peptides stymies our ability to make quantitative conclusions about proteome composition at fine-scale taxonomic resolution (e.g., because of the impact of peptide ionization efficiency or peptide cofragmentation on abundances^27^). In contrast, targeted mass spectrometry can be used to effectively measure the abundance of a single peptide with high accuracy and precision, per total protein. We combined the advantage of untargeted metaproteomics (estimating total taxon-specific protein biomass) with the advantage of targeted metaproteomics (determining concentrations of individual peptides) to estimate the number of molecules of a given peptide *per taxon-specific protein mass* (in this case, *F. kerguelensis*; Fig. 1B). To address uncertainty in both the untargeted and targeted measurements of the same sample (across technical replicates), we used a Bayesian errors-in-variables model, which carries forward error in both these types of measurements^28^.

We specifically choose to examine antioxidant proteins that were observed in the untargeted metaproteomes, and have peptides that appear to be both taxonomically- and functionally-unique for the diatom species *Fragilariopsis kerguelensis*. Many antioxidants are metal-containing, yet the contribution to total metal quotas in phytoplankton is unconstrained^29^. We used previously published genomic and transcriptomic data^30^ to confirm that 1) our peptide targets are conserved within this species, 2) these peptides differentiate between *F. cylindrus* and *F. kerguelensis*, and 3) these peptides did not spuriously map to other sequences in our database. We specifically targeted three antioxidant proteins: a copper-zinc superoxide dismutase (CuZnSOD), FeMnSOD, and peroxiredoxin. We are uncertain if the FeMnSOD contains either an Fe or Mn cofactor, but we assume that Mn is the cofactor, provide evidence below for that assumption, and hereafter refer to this protein as MnSOD. The former two proteins (CuZnSOD and MnSOD) were chosen because they have a metal cofactor and could therefore be used to assess their contribution to metal-to-carbon ratios^29^. Peroxiredoxin is an enzyme that metabolises hydrogen peroxide (without a metal cofactor), a downstream product of superoxide dismutases.

The combination of untargeted and targeted metaproteomics also allowed us to examine depth-resolved profiles of proteomic composition across stations, using the Amundsen Sea as a “natural laboratory”. Interestingly, proteomic composition across these three proteins was largely invariant with respect to depth (e.g., Station 24; Fig. 1D; all depths were <40m). Applying methods from community ecology^31^, we quantified the relationships across proteins by modelling them with a multivariate normal distribution (Materials and Methods; Supplementary Fig. S6-8), and found a weak correlation between the CuZnSOD and MnSOD (estimated Pearson correlation coefficient: 0.63; 0.1 – 0.89, 95% credible interval; Supplementary Figure S8). By identifying correlated protein abundances, our metaproteomic approach may be used in the future for identifying regulatory control mechanisms^32^ in organisms that are difficult to culture.

### Converting metaproteomic measurements to metal-to-carbon ratios

After establishing these methods for combining different metaproteomic approaches, we estimated the contribution of these proteins to elemental stoichiometry. Our approach yielded taxon-normalised estimates of protein molecules per *F. kerguelensis* protein mass (Fig. 1B), which enables simple comparisons with laboratory culture data and diatom representations in biogeochemical models. Here we focus on the metal-containing protein targets CuZnSOD and MnSOD. We estimated a median of 95 fmol MnSOD per µg *F. kerguelensis* total protein (68 – 122 fmol MnSOD per *F. kerguelensis* total protein, 95% credible interval). Assuming 3.6 pg total protein per cell^33^, this translates to ∼200,000 molecules of MnSOD per cell. Previous estimates of this protein in the brackish water diatom *Thalassiosira pseudonana* were ∼550,000 MnSOD molecules per cell^12^ (albeit *T. pseudonana* is smaller, with ∼1.4 pg total protein per cell). This comparison suggests that in both diatom species MnSOD is relatively abundant.

We calculated the contribution of MnSOD and CuZnSOD to Mn:C, Cu:C, and Zn:C using our metaproteomic measurements. First, we make the assumption that there is a 1:1 molar ratio between the peptide chosen and the protein (Fig. 2A). Second, we used previously published structural data from superoxide dismutases^34^ to identify that there is typically a single atom of a given metal per protein monomer (α parameter; Fig. 2A; for CuZnSOD we assume that there is both a single copper and a single zinc atom per SOD monomer). Third, we use previously published data^1^ on the mass ratio of protein-to-carbon in diverse phytoplankton as an informative prior estimate (λ parameter, Supplementary Fig. S9), alongside specific data on this ratio from *F. cylindrus*^33^. Finally, and most importantly, the proportion of protein metallated (ζ parameter; Fig. 2A) must be assumed to follow some probability distribution because we are not aware of data to constrain this parameter.

**Figure 2.**
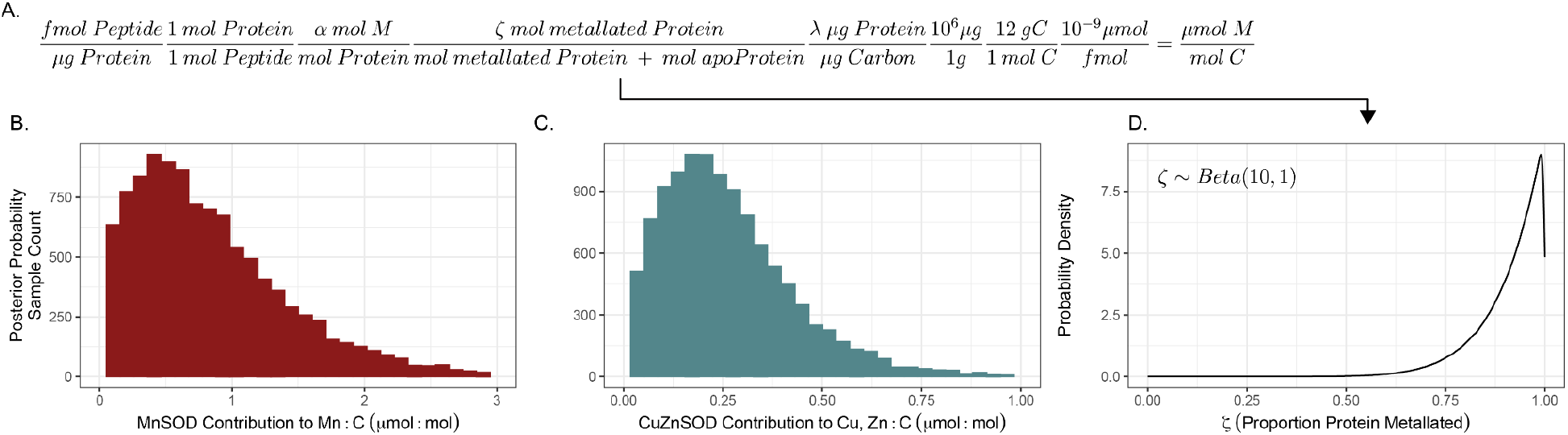
Mapping single proteins to stoichiometric ratios in natural populations of *F. kerguelensis*. (A) Calculation for converting metaproteomic measurements into the contribution to metal (*M*) to carbon ratio (described in main text). (B) Posterior probability (the probability distribution after the data have been taken into account) samples showing the contribution of MnSOD to Mn:C ratio, assuming the protein is metalated by Mn. (C) Posterior probability samples showing the contribution of CuZnSOD to Cu, Zn:C ratios. (B-C) assume that the majority of protein present is metalated following the probability density plotted in (D).

Under the assumption that the majority of superoxide dismutase is metallated (Fig. 2D; following a beta distribution with shape values 10 and 1), the contribution of Mn from MnSOD to total Mn:C in biomass has a median of 0.7 µmol Mn: mol C (0.06 – 2.2, 95% credible interval). This estimate takes into account uncertainty from both types of metaproteomic measurements, the ratio of protein to carbon, and the assumed distribution of metallated protein (see Supplementary Fig. S10 for how metallation probability distribution quantitatively impacts our estimates).

Our estimates are comparable to others that were independently derived using vastly different approaches. For example, Hawco *et al* (2018)^15^ develop a model of Mn limitation in the Southern Ocean and include a minimum Mn quota parameter representing non-photosynthetic Mn (i.e., implicitly modelling MnSOD). They prescribed a value of 1 µmol Mn: mol C, which is similar to our *in situ* measurements. We can also use the ratio of Mn in MnSOD to Mn in PSII as a comparable metric. Hawco *et al* (2022) estimated photosynthetically-derived Mn as approximately 3 µmol Mn: mol C in several species of Southern Ocean phytoplankton. Our *in situ* estimates can be divided by photosynthetically-derived Mn quotas^15^, yielding a ratio of 0.2 (for every atom of Mn in MnSOD there are 5 atoms in photosystem II). This is similar to previous Mn in MnSOD: Mn in PSII estimates of ∼0.23 from *T. pseudonana*^12^. Notably, the proportion of metallated protein follows an *assumed* probability density distribution. Intuitively, as this proportion decreases, the contribution of MnSOD to Mn:C will concomitantly decrease (Supplementary Figure S10), and therefore our field estimates could be treated as an upper bound constraint on stoichiometric contributions (just as protein-based estimates of reaction rates can provide a useful upper bound^35–37^).

How much does Mn in MnSOD contribute to the entire *F. kerguelensis* Mn quota, as a proportion of the total? To quantify this proportion, we would need *in situ* values of Mn:C for *F. kerguelensis*. We are unable to directly estimate this value with taxonomic resolution *in situ*. To address this problem, we used previously published data on Mn:C ratios^38–43^ from diatoms in the Southern Ocean to determine an empirical total quota. The median Mn:C ratio across 433 diatom measurements was 3.4 µmol Mn: mol C (see Supplementary Fig. S11 for full distribution). Therefore, our combined analyses suggest that MnSOD can account for ∼20% of the total Mn:C in this diatom species. We suspect that the remaining variation in empirical diatom Mn:C arises from Mn storage and photosynthetic proteins, which is supported by recent work using metaproteomics and measurements of particulate trace metals in the Southern Ocean^10^.

Overall, by developing an approach that combines untargeted and targeted metaproteomics, we have provided evidence that a substantial amount of Mn:C can be accounted for in MnSOD *in situ*. While it seems likely that photosynthetically-associated Mn is the dominant portion^9,15^, our results provide empirical justification for developing mechanistic models of these two individual pools of Mn, which we explore in the following section.

### Identifying the drivers of superoxide dismutases

We sought to identify the environmental drivers of superoxide dismutase abundance in an important Southern Ocean diatom, which is crucial for developing and informing mechanistic models of microbial growth^14–16^ and metal allocation. Several previous studies have modelled the relationship between electron flow and the requirement for superoxide dismutase^14,16^ based on assumptions of matching the superoxide production rate and the ability of a cell to metabolise superoxide. An overall challenge for modelling superoxide dismutases (and antioxidants in general) is, however, that they also play an important role in cell signalling^29,44–46^ and mechanistic models should account for both functions. To address this challenge, we require data on the empirical relationships between environmental conditions and superoxide dismutases for Southern Ocean phytoplankton.

We first used our *in situ* data to correlate the concentrations of dissolved Fe, dissolved Mn, temperature, and light with the *F. kerguelensis*-normalised abundances of MnSOD, CuZnSOD, and peroxiredoxin. We were unable to identify any meaningful environmental correlates (Supplementary Figure S12), which we explain with two potential reasons. First, the range of environmental conditions we observed was narrow (e.g., the dissolved iron concentrations varied from 0.06 nM to 0.31 nM; Supplementary Figure S3). Ideally we would have a wide diversity of environmental conditions, but suboptimal conditions for any taxon will drive its abundance down and therefore make measuring proteomic compositional differences more challenging. We suspect that increasingly sensitive mass spectrometry methods will mitigate this problem in the near future, facilitating the quantification of lower abundance proteins from complex mixtures more easily. Second, proteomic composition in natural assemblages is also a function of the history of environmental exposure (e.g., high iron concentrations that a cell previously was exposed to may have a lingering proteomic signature), which we were unable to quantify. We speculate that Lagrangian backwards trajectories of water parcels and environmental conditions will be a crucial tool for oceanographic expeditions that are marrying molecular and geochemical approaches in the future.

Cultures of the related Southern Ocean diatom *Fragilariopsis cylindrus* offer a controlled model for quantifying proteomic changes as a function of trace metal concentrations. We used a recently published dataset from our group to determine how dissolved Fe and Mn concentrations influence the abundance of various superoxide dismutases in *F. cylindrus*. Data-independent acquisition mass spectrometry was used to quantify over 8000 proteins from 12 culture samples. Here we examined the influence of dissolved Mn and Fe on superoxide dismutases, given the biogeochemical importance of these metals in the Southern Ocean. Iron mostly modulates the production of superoxide via impacting total photosynthetic electron flux. Previous work and observations from temperate diatoms supported the notion that MnSOD abundance should be tuned to match the supply of endogenous superoxide produced^14,16^, i.e., increased MnSOD under high iron. Surprisingly, we found that neither of the two MnSODs in *F. cylindrus* varied as a function of dissolved Fe (Fig. 3A, B), nor did the abundance of CuZnSOD (Fig. 3C). Our observations in this organism are therefore counter to the aforementioned mechanistic models, although we are unable to rule out that MnSOD activity is modulated independently of its kinetics or concentration^37^ (perhaps via a post translational modification or trafficking across cellular compartments). Quantitatively disentangling how Fe influences endogenous superoxide production (e.g., via leaky electron transport, total electron flux), and determining the consequences of varying superoxide concentrations (e.g., signalling cascades, cellular damage^29^), will be crucial for resolving the relationship between superoxide dismutases and Fe.

**Figure 3.**
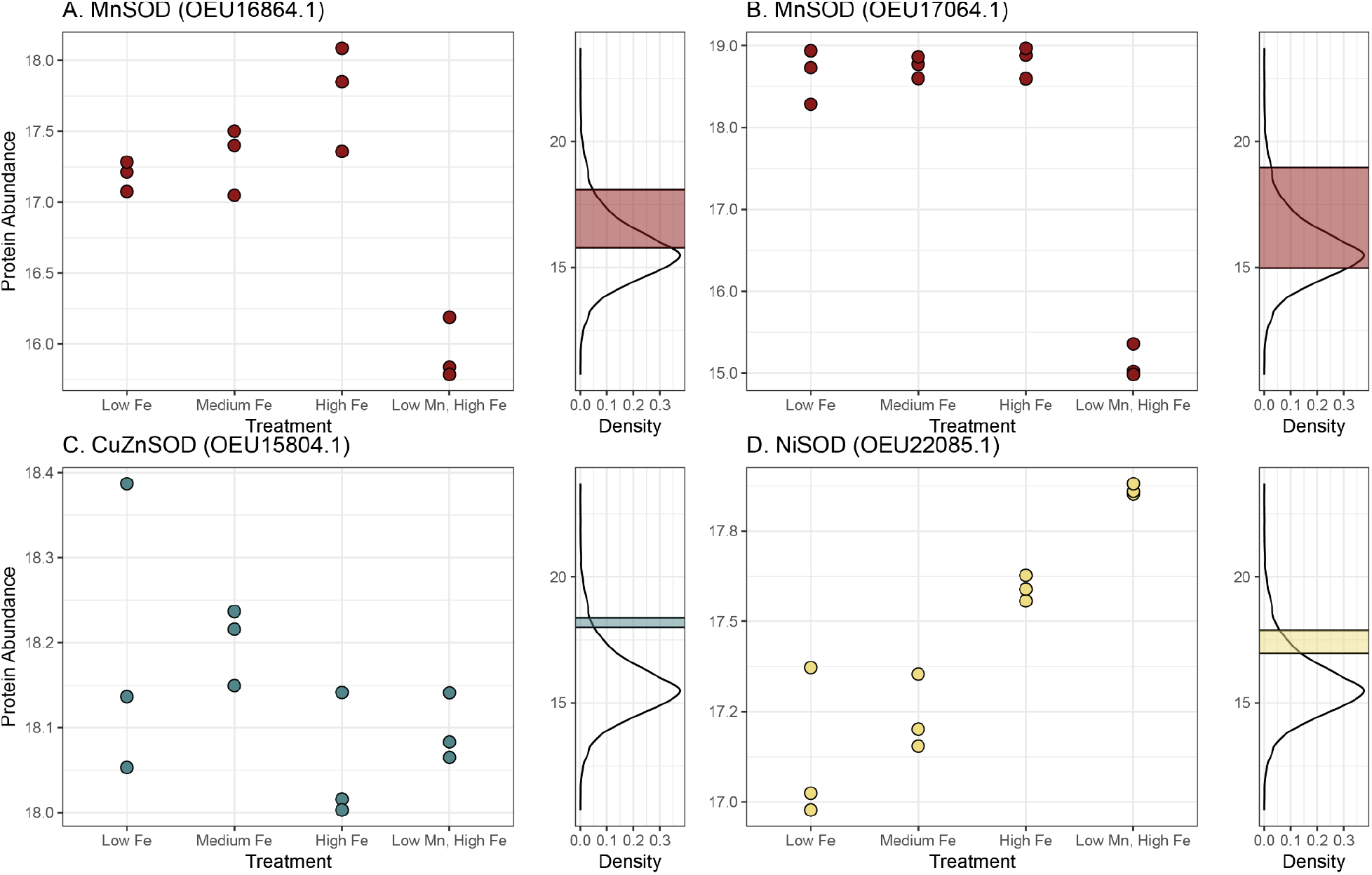
Trace-metal dependence of superoxide dismutase abundances from culture proteomics of the cold-water diatom *Fragilariopsis cylindrus*. (A-D) Superoxide dismutase protein abundances across twelve cultures of *F. cylindrus*. There were four treatments: Low Fe, Medium Fe, and High Fe (which were all supplemented with Mn). The fourth treatment was high Fe but low Mn. Density distributions of all observed proteins are plotted adjacent to the main panels to contextualize the abundances of these superoxide dismutases, with the coloured blocks showing the maximum and minimum observed protein abundance for the corresponding protein. (A-B) MnSODs; (C) CuZnSOD; and (D) NiSOD. Note that there is a fifth superoxide dismutase that we observed, a CuZnSOD, but it was only detected in a subset of conditions and was much lower in abundance.

Under high Fe and low Mn conditions, MnSOD abundance dramatically decreased. Our results indicate that low MnSOD is a biomarker for Mn limitation, though the abundance of this protein should be examined under low Fe, low Mn conditions in the future. Under Mn limiting conditions, we observed an increase in the abundance of NiSOD. This is suggestive of a compensation mechanism, where the supply of superoxide is managed via NiSOD under Mn limitation, and that NiSOD should be examined as an additional potential marker for Mn limitation. Full factorial experiments under relevant environmental conditions would be useful for disentangling the empirical drivers of superoxide dismutase abundances.

## Conclusions

Overall, we have used metaproteomes from the Southern Ocean to empirically estimate the contribution of specific proteins to metal-to-carbon ratios. Our work highlights an unknown that is necessary for further connecting metaproteomic data to metal stoichiometry: the proportion of protein metallated. Work in metalloproteomics^47,48^ on phytoplankton adapted to iron-limited regions of the ocean would be highly useful in further constraining these estimates.

Recent biogeochemical modelling efforts that introduce Mn as a limiting nutrient have done so with a parameter that represents the non-photosynthetic Mn quota (in addition to the variable photosynthetic Mn quota). We have provided a quantitative, molecular justification for this non-photosynthetic Mn quota parameter. Further, our culture proteomics data suggest that the simplest model assumption with the most empirical support is that MnSOD is constitutively expressed (i.e., represents a single constant value), or decreases under high Fe and low Mn conditions. More broadly, ecosystem models are essential tools for quantifying and predicting how organisms interact with their environments at a large scale. We have provided a path towards quantitatively integrating molecular data into ecosystem models.

## Methods

### Oceanographic Sampling of the Amundsen Sea Polynya

Samples were collected in the Amundsen Sea Polynya, Southern Ocean from December 2017 until February 2018 aboard the icebreaker RV Araon, from 15 different stations (locations with unique latitude and longitude). After the CTD was brought aboard the ship, the entire sampling system was moved into a cleanroom environment. Water was taken from the trace metal clean CTD (Seabird SBE 911+^17^) and filtration for metaproteomics began between 1 to 2.5 hours after samples were brought aboard the ship. Keeping the water containers on ice packs, we filtered water using a peristaltic pump through a series of connected polycarbonate filters of decreasing size (12.0, 3.0, and 0.2 *μ*m pore sizes) for metaproteomics. We limited filtration time to a maximum of 1.5 hours or until the filters clogged, and subsequently stored at −80 °C until protein extraction. We also collected water for analyzing dissolved and particulate trace metal concentrations at corresponding depths to the metaproteomic samples, which are discussed elsewhere^18,19^. Filters for measuring particulate metal concentrations had a pore size of 0.45 μm and were not size fractionated, which limits direct comparisons between metaproteomic measurements and particulate metal concentrations.

### Metaproteomic Sample Preparation

Proteins were extracted from only the 3.0 *μ*m filters frozen in cryovials with the following method. We focused on the 3.0 *μ*m filter size to explore *Fragilariopsis* spp. proteomic composition, which should be predominantly captured on this filter size. Protein extraction buffer (0.1 M Tris/HCl, pH 7.5, 5% glycerol, 5 mM EDTA, 2% SDS) was put into the cryovial, after which it was incubated at 95 °C for 15 minutes. Filters with extraction buffer were then sonicated on ice (15 seconds on, 15 seconds off, 2 minutes total sonication time, 50% amplitude 125 W, Qsonica Sonicator Q125, Newtown, Connecticut, USA). After sonication, we incubated the sample at room temperature for 30 minutes. Extracted protein (in buffer solution) was then removed from the cryovial, centrifuged at 15,000 G for 30 minutes to pellet cell debris, and the supernatant was removed and stored at −80 °C. We measured the total protein concentration using a BCA assay (Thermo Fisher Scientific, California, USA) to calculate the total *μ*g protein per volume seawater for this size fraction.

We then reduced and alkylated the extracted protein, and removed the SDS extraction buffer using S-traps (Protifi, Farmingdale, New York, USA). We first prepared solutions of 500 mM dithiothreitol (DTT) and 500 mM iodoacetamide (IAM) in 50 mM ammonium bicarbonate. Protein was reduced with DTT, bringing up the concentration to 5 mM and incubating at 37 °C for one hour in a Thermomixer (F1.5, Eppendorf, Hamburg, Germany) at 350 RPM. Reduced protein was then cooled to room temperature, and alkylated using IAM, bringing the concentration to three times that of DTT (15 mM). After incubating in the dark for 30 minutes at room temperature, we then quenched the reaction with 5 mM of DTT. We denatured the extracted proteins with 12% phosphoric acid by bringing it to 1.2% by volume. Our samples were then diluted with S-trap buffer (1:7, sample: S-trap buffer; 90% methanol in 100 mM triethylammonium bicarbonate, acidified to 7.1 pH with phosphoric acid). The sample and S-trap mixture were then loaded onto the S-traps, which were kept on a vacuum manifold but prevented from becoming completely dry. After the sample and S-trap mixture were fully loaded on each unit, we washed the sample with 10x 600 *μ*L of S-trap buffer to remove the SDS. For the first three washes, buffer was left on the S-trap without using the vacuum pump. S-traps with sample loaded were then centrifuged at 4000 XG for 1 minute to remove the remaining S-trap buffer.

Finally, we digested the protein using trypsin in 50 mM ammonium bicarbonate, with a ratio of 1:25 sample protein:trypsin, and incubated them at 37 °C for 16 hours. Peptides were then eluted from the S-traps with 80 *μ*L of 50 mM ammonium bicarbonate, 80 *μ*L 0.2% aqueous formic acid, and 80 *μ*L 50% acetonitrile containing 0.2% formic acid. Samples were then dried in a Vacufuge Plus (Eppendorf, Hamburg, Germany; between 3-4 hours), and then reconstituted in 3% acetonitrile and 0.1% formic acid.

The samples were desalted using 50 mg C18 columns. Columns were first conditioned with 500 *μ*L methanol, and then 500 *μ*L of 50% acetonitrile, 0.1% formic acid. Columns were then equilibrated with two aliquots of 500 *μ*L of 0.1% trifluoroacetic acid. We then increased the volume of samples that were previously reconstituted with 3% acetonitrile and 1% formic acid by adding 100 *μ*L of this solution, and then loaded the diluted sample onto the equilibrated column. Samples were then pushed through the column using a syringe, and then washed three times with 1 mL of 0.1% TFA each time, removing the salt and retaining the peptides on the column. Finally, peptides were eluted into a low binding plastic microcentrifuge tube (Thermo Fisher Scientific, California, USA) with two aliquots of 200 *μ*L of 50% acetonitrile and 0.1% formic acid, and then one aliquot of 70% acetonitrile and 0.1% formic acid. Samples were then dried down using a Vacufuge Plus (Eppendorf, Hamburg, Germany; between 5-6 hours), until only the dried peptides remained.

### Liquid chromatography mass spectrometry for untargeted metaproteomics

We used both untargeted and targeted mass spectrometry-based metaproteomics. For untargeted mass spectrometry, we used liquid chromatographic (LC) separation of the complex peptide mixture to reduce the sample complexity prior to injecting into the mass spectrometer. The LC was coupled directly to a Q Exactive hybrid quadrupole-Orbitrap mass spectrometer (Thermo Fisher Scientific, California, USA), and the entire run lasted 125 minutes, using a non-linear gradient. We used a data-dependent acquisition mass spectrometry approach with a TopN value of 8. The MS1 scans were run at 140,000 resolution, with a scan range from 400 to 2000 *m/z* and an automatic gain control target of 3E6. For the MS2 scans, we chose a resolution of 17,500, an automatic gain control target of 1E6, an isolation window of 2 *m/z*, and a scan range of 200 to 2000 *m/z*. All samples were injected into the mass spectrometer twice, with the exception of two samples that had failed injections because of challenges with the column.

### Untargeted metaproteomic bioinformatics

Metaproteomics requires a database of potential proteins to search mass spectra against. We used a custom database of metatranscriptomic sequences from the neighbouring Ross Sea, and appended those protein sequences with sequences from metagenome-assembled genomes (MAGs). We were interested in the diversity of antioxidant proteins that phytoplankton are using, so we identified antioxidant-proteins of interest from this large collection of eukaryotic MAGs^21^ using Enzyme Commission numbers associated with their sequence annotations (E.C. numbers: 1.15.1.1; 1.11.1.11; 1.11.1.5; 1.11.1.9; 1.11.1.15; 1.11.1.6). We then reduced the database size by combining protein sequences that are 95% or higher sequence similarity with CD-HIT^49^. Finally, we appended a database of common contaminants (Global Proteome Machine Organization common Repository of Adventitious Proteins). In total, there were 414498 protein sequences in our database. We then used MSGF+^50^ within OpenMS^51,52^ with the following settings: fixed cysteine carbamidomethyl, and variable methionine oxidation, N-terminal glutamate to pyroglutamate, deamidated asparagine, and deamidated glutamine. A 1% false discovery rate was applied at the peptide spectrum match level.

Peptides were quantified at the MS1 level with their corresponding ion intensities^53,54^. FeatureFinderIdentification is an approach that cross-maps identified MS2 spectra to unidentified features across samples. This approach requires grouping samples, and we conservatively only grouped samples that were technical replicates. Note that for two samples, we were unable to acquire duplicate injections (sample ID number 74 and 197).

The choice of database can have important consequences for estimating peptide abundances, particularly across diverse samples. We previously put forward a metric to evaluate if database dependence was variable across samples by correlating the total ion current for each sample with the sum of identified peptide abundances^25^. If this correlation is high, that is evidence database dependence will not impact estimated peptide abundances. In these samples, we found a group of samples that did not correlate well with total ion current. We therefore used a database-independent normalization: we predicted peptide-like features using Dinosaur^55^, and then used the sum of peptide-like features as a normalization constant. Each raw peptide intensity was normalized by the sum of peptide-like features.

### Targeted metaproteomics liquid chromatography and mass spectrometry

We chose peptides to measure several antioxidant proteins in *F. kerguelensis* in the Amundsen Sea samples. We restricted our peptide selection to peptides that were identified in the untargeted metaproteomes and chose peptides with the following amino acid sequences (isotopically heavy residue is bolded): VSENFAYV**V**K (derived from Fe, MnSOD); SGVCQLVYQE**L**AK (derived from peroxiredoxin); and DGSSSCGPIFNP**F**GK (derived from CuZnSOD). We further examined the taxonomic redundancy and resolution of these peptides by first ensuring that they only taxonomically map to *F. kerguelensis* within our database, and not to another group of organisms. We then determined the associated protein sequences present in *F. cylindrus*^56^ and *F. kerguelensis*^30,57^, aligned these sequences using MAFFT^58^, and the manually examined the alignments to ensure that our peptide choices map only to *F. kerguelensis* and not to *F. cylindrus*. We also ensured that these peptides map to different strains of *F. kerguelensis*^30,57^.

Targeted mass spectrometry was performed using a Dionex Ultimate 3000 UPLC system interfaced to a TSQ Quantiva triple-stage quadrupole mass spectrometer (MS) (Thermo Scientific, Waltham, MA), fitted with a heated, low flow capillary ESI probe (HESI-II). Total protein injected per sample was 0.8 ug for most samples. In cases where less material was available 0.4 or 0.6 ug protein was injected. Isotopically labeled, heavy internal standard versions of each targeted peptide were synthesized by Thermo Scientific™ at >95% purity.

Heavy peptide standard was added to samples to yield 20 fmol on column for every sample injection. Details of mass spectrometer conditions are reported in supplement methods. Each sample was analyzed via triplicate injections using the transition list of peptides that is found in Supplemental Table 1.

### Estimating *Fragilariopsis* spp. and *F. kerguelensis* biomass

We estimated the proportion of protein attributed to *Fragilariopsis* spp. First, we identified all peptides that are taxonomically informative for a range of taxonomic groups, mostly at the phylum or genus levels. The sum of peptide intensities attributed to a specific organism can be used as a metric of relative abundance, where each individual peptide intensity is normalised to the sum of peptide-like features (described above). We wanted to consider the absolute abundance of these taxonomic groups, so we had to account for taxonomically uninformative peptides. For example, in a metaproteome of two highly similar species of equal abundances, abundances of taxonomically-informative peptides need to be rescaled so that the abundance estimates are not spuriously low. We therefore rescaled the taxonomically-informative peptide abundances such that the sum of normalised peptide abundances remained constant.

We had sufficient peptides (>50) to estimate the proportion of total protein biomass that is attributed to *Fragilariopsis* spp., however we did not have sufficient peptides to examine specific species across all samples. We therefore conservatively focused only on regions of high *Fragilariopsis* spp. biomass (stations 50, 52, 53, 57, 55, and 24). Within these stations, we multiplied the proportion of *Fragilariopsis* spp. biomass by another factor: the fraction of species-specific peptides for *F. kerguelensis*. This fraction was determined by first examining all peptides observed from the metaproteomes, and labelling those that were *Fragilariopsis* spp.-specific as either *F. cylindrus*-specific, *F. kerguelensis*-specific, or *Fragilariopsis*-genus specific. Our fraction is then the sum of *F. kerguelensis-*specific peptides divided by the sum of both *F. kerguelensis*-specific and *F. cylindrus*-specific peptides. This results in a single value that represents the proportion of total protein attributed to *F. kerguelensis*.

### Statistical modelling

We used a Bayesian statistical model to describe 1) measurement error from untargeted and targeted metaproteomics, 2) the relationship among proteins, 3) the dependence of proteins on environmental correlates, 4) the impact of single protein measurements on elemental stoichiometry. The advantage of this modelling framework is that various uncertainties are represented in a single probabilistic model. In total we had 42 samples for both types of measurements, however, *F. kerguelensis* was abundant in only a subset (14 samples had >50 *F. kerguelensis*-specific peptides). We therefore focused on this subset for fitting our statistical model. Measurement error was modelled using an errors-in-variables approach,

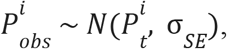

where replicate measurements are 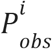, and we estimated the true underlying mean value 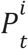 with information from the technical replicates (via the standard error, *σ*_*SE*_), following a normal distribution (*N*). Superscript *i* refers to either the untargeted or targeted measurements (*i* ∈ {*U, T*}). For the targeted measurements, we typically had 3 replicate injections (with a range of 2–8 injections). For the untargeted measurements, there were mostly two replicate injections (samples ID 74 and 197 only had one injection; for these samples we used the average standard error). Given the untargeted measurements result in a value constrained by 0 and 1, we used a truncated normal distribution and a weakly informative prior with a mean of 0.1 and standard deviation of 0.5.

We transformed the two errors-in-variables quantities into a ratio, therefore arriving at the desired quantity, fmol Peptide per *F. kerguelensis* protein (i.e., 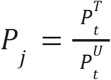 where superscript *j* refers to the specific protein targeted). Uncertainty from the measurement is therefore incorporated from both the numerator and denominator.

We modelled taxon-normalized peptide abundances (*P*_*j*_) as a multivariate normal distribution where the mean values are a function of environmental correlates. This approach was borrowed from community ecology^31^, where there is both an environmental and biotic dependence on a species’ abundance. From a molecular perspective, this can be interpreted as the influence of an environmental trigger (e.g., promoter activity dependent on an environmental variable) as well as two proteins having a shared regulator. Specifically, we modelled the estimated protein abundances as

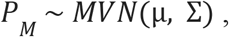

where *P*_*M*_ is a matrix of estimated, taxon-normalised peptide abundances (with 3 columns and 14 rows), which are assumed to follow a multivariate normal distribution (*MVN*), where µ is a 3-element vector of mean abundances, and Σ is a 3 × 3 covariance matrix.

Lastly, we wanted to explain variation in the abundance of each protein, so we examined how the abundance varied as a function of several environmental covariates. Specifically, we examined dissolved Fe and Mn concentrations, light, and temperature. These measurements were taken at similar depths (see Supplementary Figures S2, S3). We therefore set the mean abundance as a function of environmental correlates,

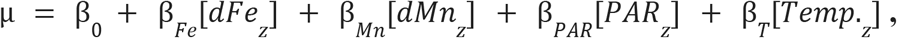

where β_0_ is a three-element vector of intercept coefficients, β_*Fe*_ represents the set of coefficients for dissolved Fe, etc. Prior to fitting this model, we z-score transformed each environmental covariate.

We used the fitted statistical model to make posterior predictive distributions for connecting the metaproteomic measurements to units of elemental stoichiometry, in particular metal-to-carbon ratios. We did not observe any meaningful environmental covariates (most estimated β coefficient values had large uncertainties and overlapped with zero, except β_0_). We therefore used the intercept value (β_0_) for generating posterior predictive distributions. We assumed a 1:1 molar ratio of the peptide to protein (α in Fig. 2A). We assumed various beta distributions, as discussed in the main text, for the proportion of metallated protein (ζ in Fig. 2A). Finally, we empirically estimated the mass ratio of protein to carbon (λ in Fig. 2A). First, we used informative priors from a range of several phytoplankton^1^, with a mean of 0.62 and a standard deviation of 0.34,

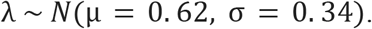

We then estimated λ using previously published data from *F. cylindrus*,

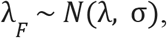

where σ is the standard deviation. With these estimated parameters (λ and σ), we generated posterior predictions for the mass ratio of protein to carbon for the final calculation. The model was fitted using Stan and Rstan^59,60^.

## Data Availability

Proteomics data will be deposited in PRIDE upon publication. Oceanographic data and other relevant data sources are available in Zenodo: https://doi.org/10.5281/zenodo.17194677.

## Code

All code for the analyses and figures is available at https://github.com/jspmccain/amundsen-prot.

## Acknowledgements

We are grateful to the crew of the RV Araon and all scientists aboard, specifically the trace metal sampling team Mathijs Van Manen, Sven Pont, Hung-An Tian, and Charlotte Eich for their help with sample collection. We are also grateful to Alexander Cohen and Christopher Hughes in the Biological Mass Spectrometry Core Facility at Dalhousie University. Gregory Britten, Bob Carpenter, Eric Novik, and Jacqueline Buros Novik provided helpful discussions about the Bayesian statistical model. Nathalie Joli assisted with the raw data on *F. cylindrus* protein and carbon quotas. Thank you to Benjamin Twining for assistance in aggregating the single cell data on diatom Mn:C. J.S.P.M. is currently a Damon Runyon Fellow supported by the Damon Runyon Cancer Research Foundation (DRG-2470-22), and was previously supported by an NSERC CGS-D and a Transatlantic Ocean System Science and Technology Fellowship. This work was part of the FePhyrus project (ALWPP.2016.020), which was supported by the Netherlands Polar Programme (NPP), with financial aid from the Dutch Research Council (NWO). The ANA08B expedition was supported by the Korea Polar Research Institute, grant KOPRI PE25110. This work was also supported by NSERC Discovery Grant RGPIN-2015-05009 to EMB, Simons Foundation Grant 504183 to EMB, Simons Foundation CBIOMES Award ID 1001702 to EMB, Canada Research Chair Support to EMB.

## References

1. Liefer, J. D. et al. The Macromolecular Basis of Phytoplankton C:N:P Under Nitrogen Starvation. Front. Microbiol. 10, 763 (2019).

2. Inomura, K., Deutsch, C., Jahn, O., Dutkiewicz, S. & Follows, M. J. Global patterns in marine organic matter stoichiometry driven by phytoplankton ecophysiology. Nat. Geosci. 15, 1034–1040 (2022).

3. Tagliabue, A. et al. An iron cycle cascade governs the response of equatorial Pacific ecosystems to climate change. Global Change Biology 26, 6168–6179 (2020).

4. Buma, A. G. J., De Baar, H. J. W., Nolting, R. F. & Van Bennekom, A. J. Metal enrichment experiments in the Weddell-Scotia Seas: Effects of iron and manganese on various plankton communities. Limnol. Oceanogr. 36, 1865–1878 (1991).

5. Wu, M. et al. Manganese and iron deficiency in Southern Ocean Phaeocystis antarctica populations revealed through taxon-specific protein indicators. Nat Commun 10, 3582 (2019).

6. Middag, R., De Baar, H. J. W., Klunder, M. B. & Laan, P. Fluxes of dissolved aluminum and manganese to the Weddell Sea and indications for manganese co‐limitation. Limnology & Oceanography 58, 287–300 (2013).

7. Browning, T. J., Achterberg, E. P., Engel, A. & Mawji, E. Manganese co-limitation of phytoplankton growth and major nutrient drawdown in the Southern Ocean. Nat Commun 12, 884 (2021).

8. Browning, T. J. et al. Strong responses of Southern Ocean phytoplankton communities to volcanic ash. Geophysical Research Letters 41, 2851–2857 (2014).

9. Raven, J. A. Predictions of Mn and Fe use efficiencies of phototrophic growth as a function of light availability for growth and of C assimilation pathway. New Phytologist 116, 1–18 (1990).

10. Jabre, L. J. et al. Elemental allocation to molecular drivers of biogeochemistry in the Southern Ocean. In prep.

11. Dupont, C. L., Neupane, K., Shearer, J. & Palenik, B. Diversity, function and evolution of genes coding for putative Ni‐containing superoxide dismutases. Environmental Microbiology 10, 1831–1843 (2008).

12. Wolfe-Simon, F., Starovoytov, V., Reinfelder, J. R., Schofield, O. & Falkowski, P. G. Localization and Role of Manganese Superoxide Dismutase in a Marine Diatom. Plant Physiol. 142, 1701–1709 (2006).

13. Middag, R., De Baar, H. J. W., Laan, P., Cai, P. H. & Van Ooijen, J. C. Dissolved manganese in the Atlantic sector of the Southern Ocean. Deep Sea Research Part II: Topical Studies in Oceanography 58, 2661–2677 (2011).

14. Anugerahanti, P. & Tagliabue, A. Process controlling iron–manganese regulation of the Southern Ocean biological carbon pump. Phil. Trans. R. Soc. A. 381, 20220065 (2023).

15. Hawco, N. J., Tagliabue, A. & Twining, B. S. Manganese Limitation of Phytoplankton Physiology and Productivity in the Southern Ocean. Global Biogeochemical Cycles 36, e2022GB007382 (2022).

16. McCain, J. S. P. et al. Cellular costs underpin micronutrient limitation in phytoplankton. Sci. Adv. 7, eabg6501 (2021).

17. De Baar, H. J. W. et al. Titan: A new facility for ultraclean sampling of trace elements and isotopes in the deep oceans in the international Geotraces program. Marine Chemistry 111, 4–21 (2008).

18. Tian, H.-A. et al. The biogeochemistry of zinc and cadmium in the Amundsen Sea, coastal Antarctica. Marine Chemistry 249, 104223 (2023).

19. Van Manen, M. et al. The role of the Dotson Ice Shelf and Circumpolar Deep Water as driver and source of dissolved and particulate iron and manganese in the Amundsen Sea polynya, Southern Ocean. Marine Chemistry 246, 104161 (2022).

20. Jabre, L. J. et al. Molecular underpinnings and biogeochemical consequences of enhanced diatom growth in a warming Southern Ocean. Proc. Natl. Acad. Sci. U.S.A. 118, e2107238118 (2021).

21. Delmont, T. O. et al. Functional repertoire convergence of distantly related eukaryotic plankton lineages abundant in the sunlit ocean. Cell Genomics 2, 100123 (2022).

22. Eich, C. et al. Ecological Importance of Viral Lysis as a Loss Factor of Phytoplankton in the Amundsen Sea. Microorganisms 10, 1967 (2022).

23. Smith, V. H. Low Nitrogen to Phosphorus Ratios Favor Dominance by Blue-Green Algae in Lake Phytoplankton. Science 221, 669–671 (1983).

24. Cefarelli, A. O. et al. Diversity of the diatom genus Fragilariopsis in the Argentine Sea and Antarctic waters: morphology, distribution and abundance. Polar Biol 33, 1463–1484 (2010).

25. McCain, J. S. P., Allen, A. E. & Bertrand, E. M. Proteomic traits vary across taxa in a coastal Antarctic phytoplankton bloom. ISME J 16, 569–579 (2022).

26. Kleiner, M. et al. Assessing species biomass contributions in microbial communities via metaproteomics. Nat Commun 8, 1558 (2017).

27. McCain, J. S. P. & Bertrand, E. M. Prediction and Consequences of Cofragmentation in Metaproteomics. J. Proteome Res. 18, 3555–3566 (2019).

28. Britten, G. L. et al. Evaluating the Benefits of Bayesian Hierarchical Methods for Analyzing Heterogeneous Environmental Datasets: A Case Study of Marine Organic Carbon Fluxes. Front. Environ. Sci. 9, 491636 (2021).

29. McCain, J. S. P. & Bertrand, E. M. Phytoplankton antioxidant systems and their contributions to cellular elemental stoichiometry. Limnol Oceanogr Letters 7, 96–111 (2022).

30. Keeling, P. J. et al. The Marine Microbial Eukaryote Transcriptome Sequencing Project (MMETSP): Illuminating the Functional Diversity of Eukaryotic Life in the Oceans through Transcriptome Sequencing. PLoS Biol 12, e1001889 (2014).

31. Tikhonov, G., Abrego, N., Dunson, D. & Ovaskainen, O. Using joint species distribution models for evaluating how species‐to‐species associations depend on the environmental context. Methods Ecol Evol 8, 443–452 (2017).

32. Nicolas, P. et al. Condition-Dependent Transcriptome Reveals High-Level Regulatory Architecture in Bacillus subtilis. Science 335, 1103–1106 (2012).

33. Joli, N. et al. Hypometabolism to survive the long polar night and subsequent successful return to light in the diatom Fragilariopsis cylindrus. New Phytologist 241, 2193–2208 (2024).

34. Sheng, Y. et al. Superoxide Dismutases and Superoxide Reductases. Chem. Rev. 114, 3854–3918 (2014).

35. Saito, M. A. et al. Abundant nitrite-oxidizing metalloenzymes in the mesopelagic zone of the tropical Pacific Ocean. Nat. Geosci. 13, 355–362 (2020).

36. Roberts, M. E. et al. Rubisco in high Arctic tidewater glacier‐marine systems: A new window into phytoplankton dynamics. Limnology & Oceanography lno.12525 (2024) doi:10.1002/lno.12525.

37. McCain, JSP, Britten, G, Hackett, SR, Follows, MJ, & Li, G-W. Microbial reaction rate estimation using proteins and proteomes. The ISME Journal (2025).

38. Sofen, L. E. et al. Authigenic Iron Is a Significant Component of Oceanic Labile Particulate Iron Inventories. Global Biogeochemical Cycles 37, e2023GB007837 (2023).

39. Twining, B. S. et al. Metal quotas of plankton in the equatorial Pacific Ocean. Deep Sea Research Part II: Topical Studies in Oceanography 58, 325–341 (2011).

40. Twining, B. S. et al. Taxonomic and nutrient controls on phytoplankton iron quotas in the ocean. Limnol Oceanogr Letters 6, 96–106 (2021).

41. Twining, B. S., Nuñez-Milland, D., Vogt, S., Johnson, R. S. & Sedwick, P. N. Variations in Synechococcus cell quotas of phosphorus, sulfur, manganese, iron, nickel, and zinc within mesoscale eddies in the Sargasso Sea. Limnology & Oceanography 55, 492–506 (2010).

42. Sofen, L. E. et al. Trace metal contents of autotrophic flagellates from contrasting open‐ocean ecosystems. Limnol Oceanogr Letters 7, 354–362 (2022).

43. Twining, B. S., Baines, S. B. & Fisher, N. S. Element stoichiometries of individual plankton cells collected during the Southern Ocean Iron Experiment (SOFeX). Limnol. Oceanogr. 49, 2115–2128 (2004).

44. Mittler, R. ROS Are Good. Trends in Plant Science 22, 11–19 (2017).

45. Mittler, R. et al. ROS signaling: the new wave? Trends in Plant Science 16, 300–309 (2011).

46. Case, A. On the Origin of Superoxide Dismutase: An Evolutionary Perspective of Superoxide-Mediated Redox Signaling. Antioxidants 6, 82 (2017).

47. Mazzotta, M. G., McIlvin, M. R. & Saito, M. A. Characterization of the Fe metalloproteome of a ubiquitous marine heterotroph, Pseudoalteromonas (BB2-AT2): multiple bacterioferritin copies enable significant Fe storage. Metallomics 12, 654–667 (2020).

48. Young, T. R. et al. Calculating metalation in cells reveals CobW acquires CoII for vitamin B12 biosynthesis while related proteins prefer ZnII. Nat Commun 12, 1195 (2021).

49. Fu, L., Niu, B., Zhu, Z., Wu, S. & Li, W. CD-HIT: accelerated for clustering the next-generation sequencing data. Bioinformatics 28, 3150–3152 (2012).

50. Kim, S. & Pevzner, P. A. MS-GF+ makes progress towards a universal database search tool for proteomics. Nat Commun 5, 5277 (2014).

51. Röst, H. L., Schmitt, U., Aebersold, R. & Malmström, L. pyOpenMS: A Python‐based interface to the OpenMS mass‐spectrometry algorithm library. Proteomics 14, 74–77 (2014).

52. Röst, H. L. et al. OpenMS: a flexible open-source software platform for mass spectrometry data analysis. Nat Methods 13, 741–748 (2016).

53. Weisser, H. & Choudhary, J. S. Targeted Feature Detection for Data-Dependent Shotgun Proteomics. J. Proteome Res. 16, 2964–2974 (2017).

54. Weisser, H. et al. An Automated Pipeline for High-Throughput Label-Free Quantitative Proteomics. J. Proteome Res. 12, 1628–1644 (2013).

55. Teleman, J., Chawade, A., Sandin, M., Levander, F. & Malmström, J. Dinosaur: A Refined Open-Source Peptide MS Feature Detector. J. Proteome Res. 15, 2143–2151 (2016).

56. Mock, T. et al. Evolutionary genomics of the cold-adapted diatom Fragilariopsis cylindrus. Nature 541, 536–540 (2017).

57. Johnson, L. K., Alexander, H. & Brown, C. T. Re-assembly, quality evaluation, and annotation of 678 microbial eukaryotic reference transcriptomes. GigaScience 8, giy158 (2019).

58. Katoh, K. & Standley, D. M. MAFFT Multiple Sequence Alignment Software Version 7: Improvements in Performance and Usability. Molecular Biology and Evolution 30, 772–780 (2013).

59. Stan Development Team. Stan Modeling Language Users Guide and Reference Manual, 2.34. https://mc-stan.org. (2024).

60. Stan Development Team. “RStan: the R interface to Stan.” R package version 2.32.5, https://mc-stan.org/. (2024).

