## Supplementary Materials for "Superoxide dismutases shape manganese stoichiometry in Southern Ocean diatoms"

### Supplementary Methods

#### Targeted metaproteomics liquid chromatography and mass spectrometry

Targeted metaproteomic analysis was performed using a Dionex Ultimate 3000 UPLC coupled with a TSQ Quantiva mass spectrometer (MS). The MS was equipped with a heated low flow capillary ESI probe (HESI-II) with the following settings: spray voltage of 3500 V, sheathgas 5, auxillary gas 2, ion transfer tube temperature of 325 °C, vapor gas 70 °C and Chrom filter setting of 10 s. Samples were diluted with 1% FA, 3% ACN to a final peptide concentration of approximately 0.167 µg/µL. Each sample was spiked with 3.3 fmol/µl of each heavy isotope-labelled internal standard for each peptide (Thermo Scientific, Supplemental Data Table 1). 6 µl injections were performed in triplicate. Samples were concentrated on an Acclaim Pepmap C18 loading column (0.3 x 5 mm) and separated with an Acclaim Pepmap C18 analytical column (0.3 x 150 mm, 2 µm particle size, 100 Å) at a flow rate of 5 ul/min with a linear gradient from 4% B solvent to 45% B solvent over 40 minutes; the total LC/MS run time including washing and re-equilibration was 61 minutes. Solvent A was 0.1 % formic acid in HPLC grade water and solvent B was 80% acetonitrile, 0.1% formic acid in HPLC grade water. Selected reaction monitoring (SRM) transitions were optimized on our instrument using the Quantiva transition optimization tool. The method contained 244 transitions for 52 peptides (Supplementary Data Table S2) although only three peptides are discussed here (Supplementary Data Table S1). The transition dwell time was 5 msec, Q1 and Q3 resolution was set to 0.7 (FWHM), automatically calibrated RF lens setting, and a collision gas pressure of 2.5 mTorr. All raw targeted metaproteomic data obtained from the mass spectrometer was processed using Skyline-daily software[1]. Peptide concentrations were calculated from the mass spectrometry data by multiplying the peak area of each peptide of interest by the ratio of moles of the heavy isotope-labeled version of that peptide added to the peak area corresponding to that heavy isotope labeled peptide.

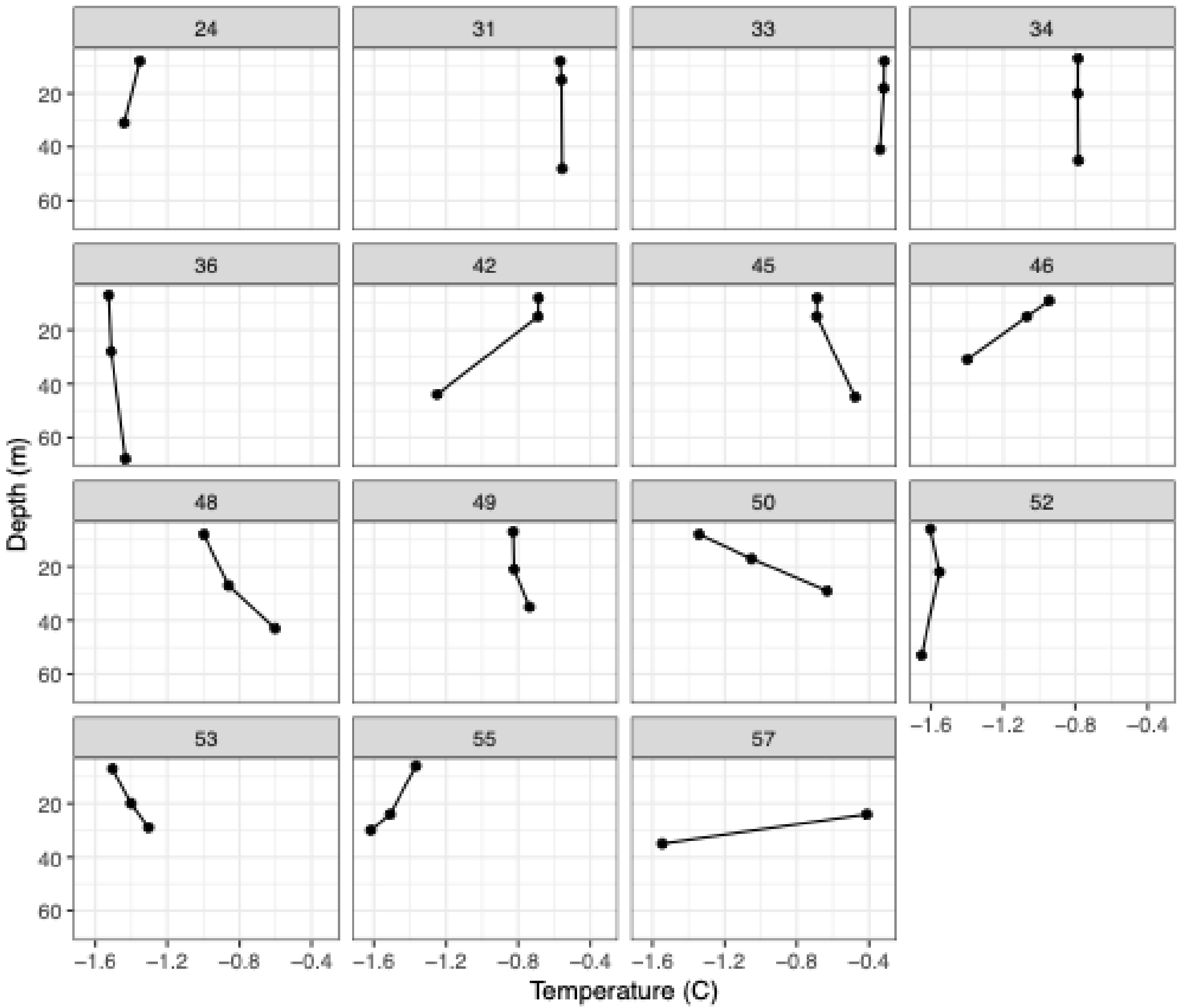


**Supplementary Figure S1.** Depth profiles of temperature across stations in the Amundsen Sea Polynya. Each panel refers to a unique station (shown in Fig. 1 in main text).

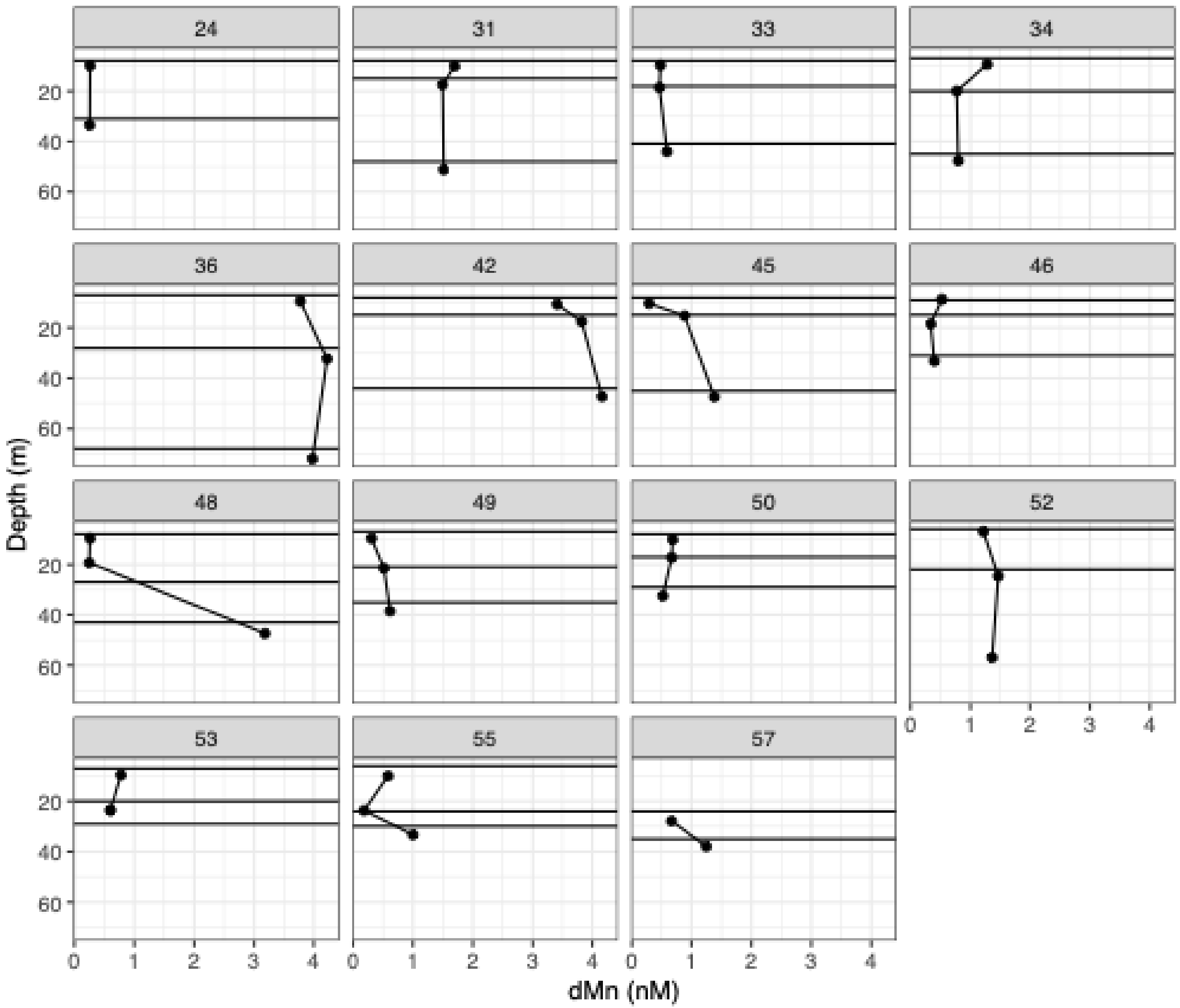


**Supplementary Figure S2**. Depth profiles of dissolved Mn across stations in the Amundsen Sea Polynya. Trace metals were sampled with a different bottle, and so the depth of the protein samples were slightly different than the trace metal concentration measurements. Protein bottles (the water we used for metaproteomics) are shown as horizontal lines. Each panel refers to a unique station (shown in Fig. 1 in main text).

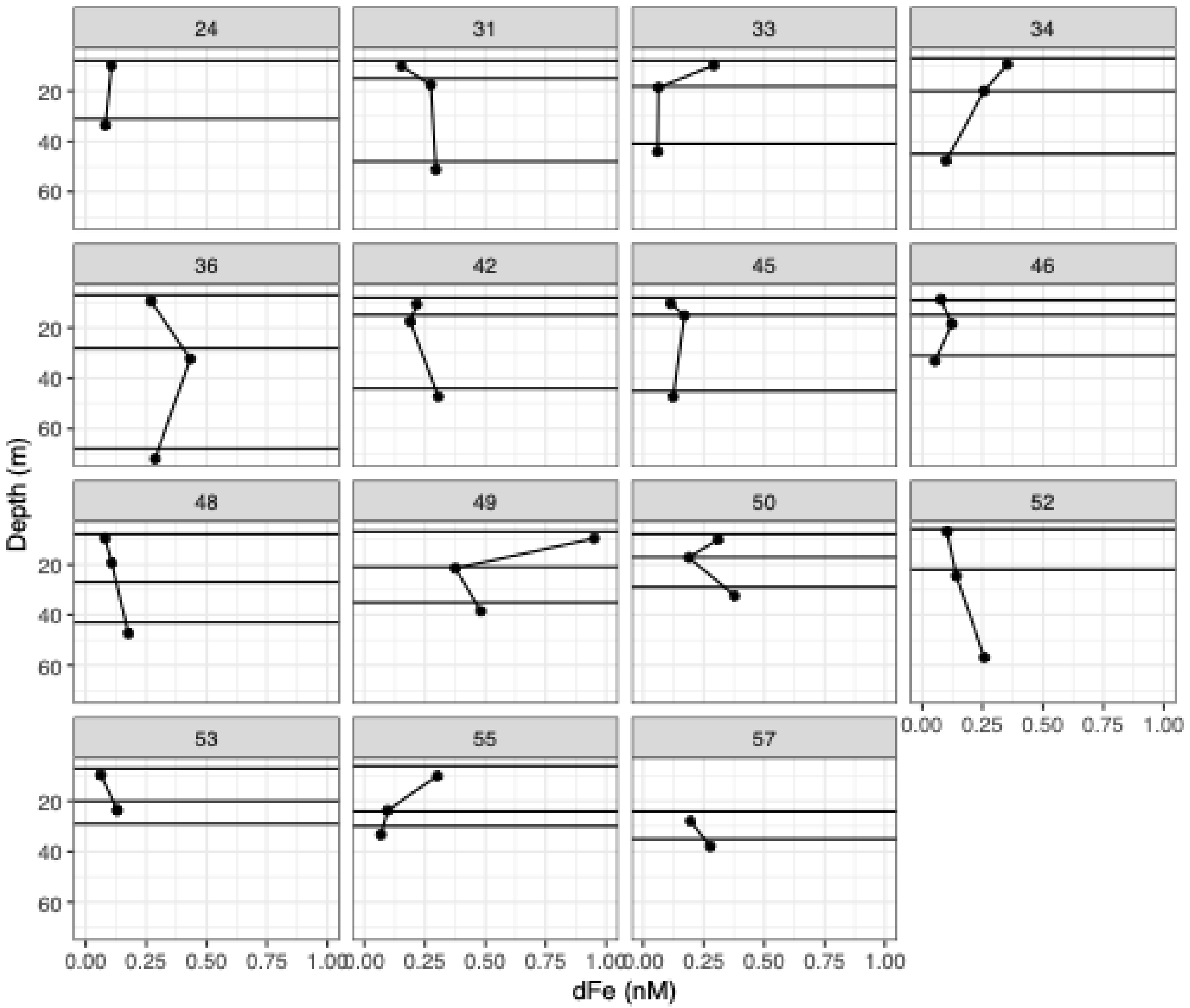


**Supplementary Figure S3.** Depth profiles of dissolved Fe across stations in the Amundsen Sea Polynya. Trace metals were sampled with a different bottle, and so the depth of the protein samples were slightly different than the trace metal concentration measurements. Protein bottles (the water we used for metaproteomics) are shown as horizontal lines. Each panel refers to a unique station (shown in Fig. 1 in main text).

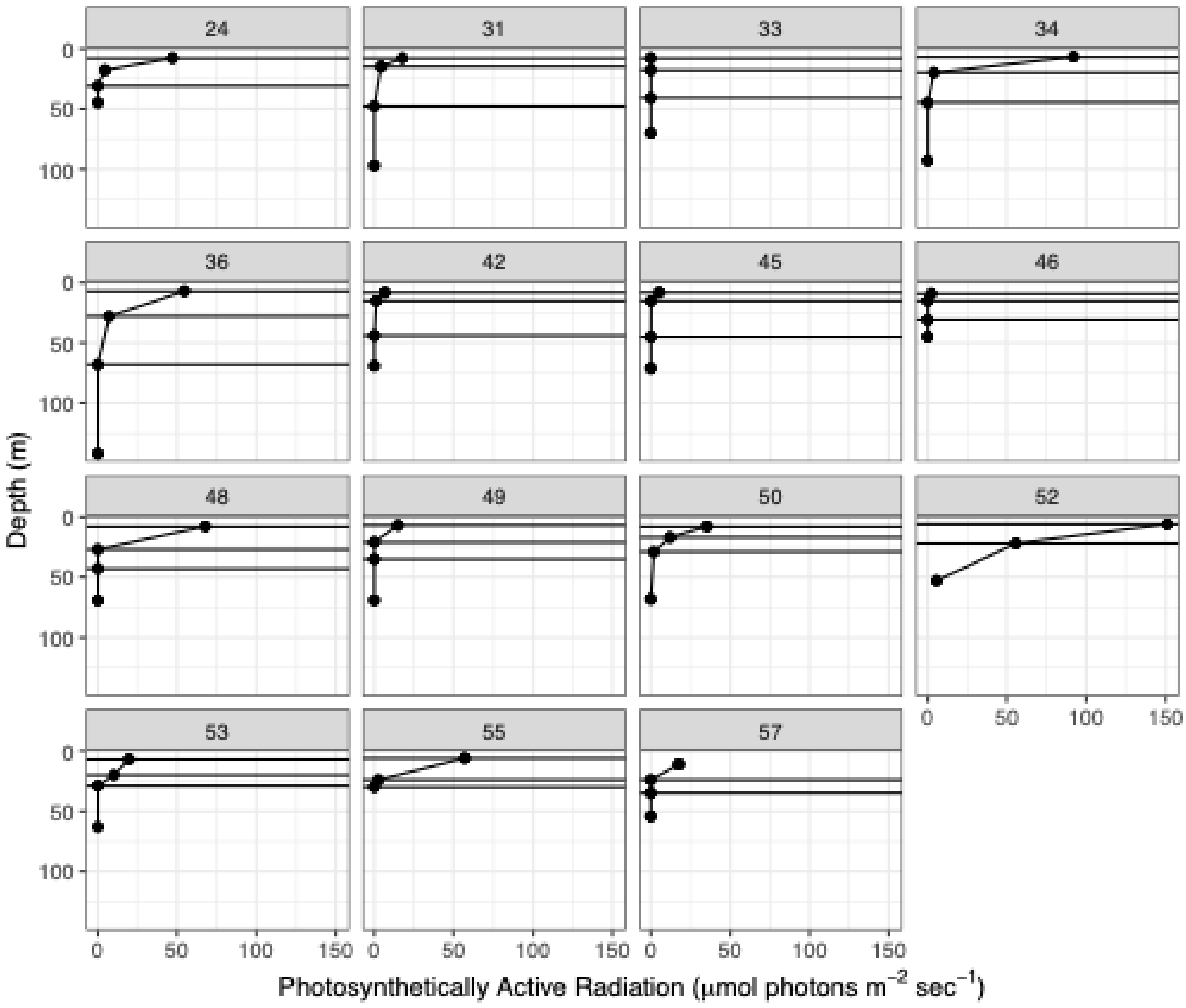


**Supplementary Figure S4.** Depth profiles of photosynthetically active radiation across stations in the Amundsen Sea Polynya. Horizontal lines shown the bottles that were used for metaproteomics. Each panel refers to a unique station (shown in Fig. 1 in main text).

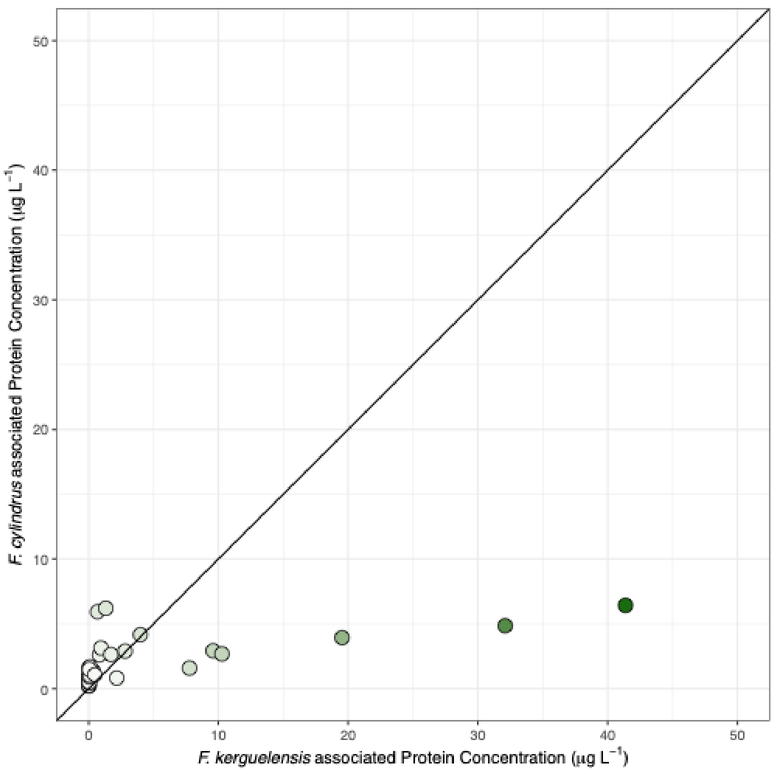


**Supplementary Figure S5**. Abundance estimates of *F. kerguelensis* protein compared to *F. cylindrus*. Colour refers to the estimated concentration of all diatom-associated protein (as shown in the main text).

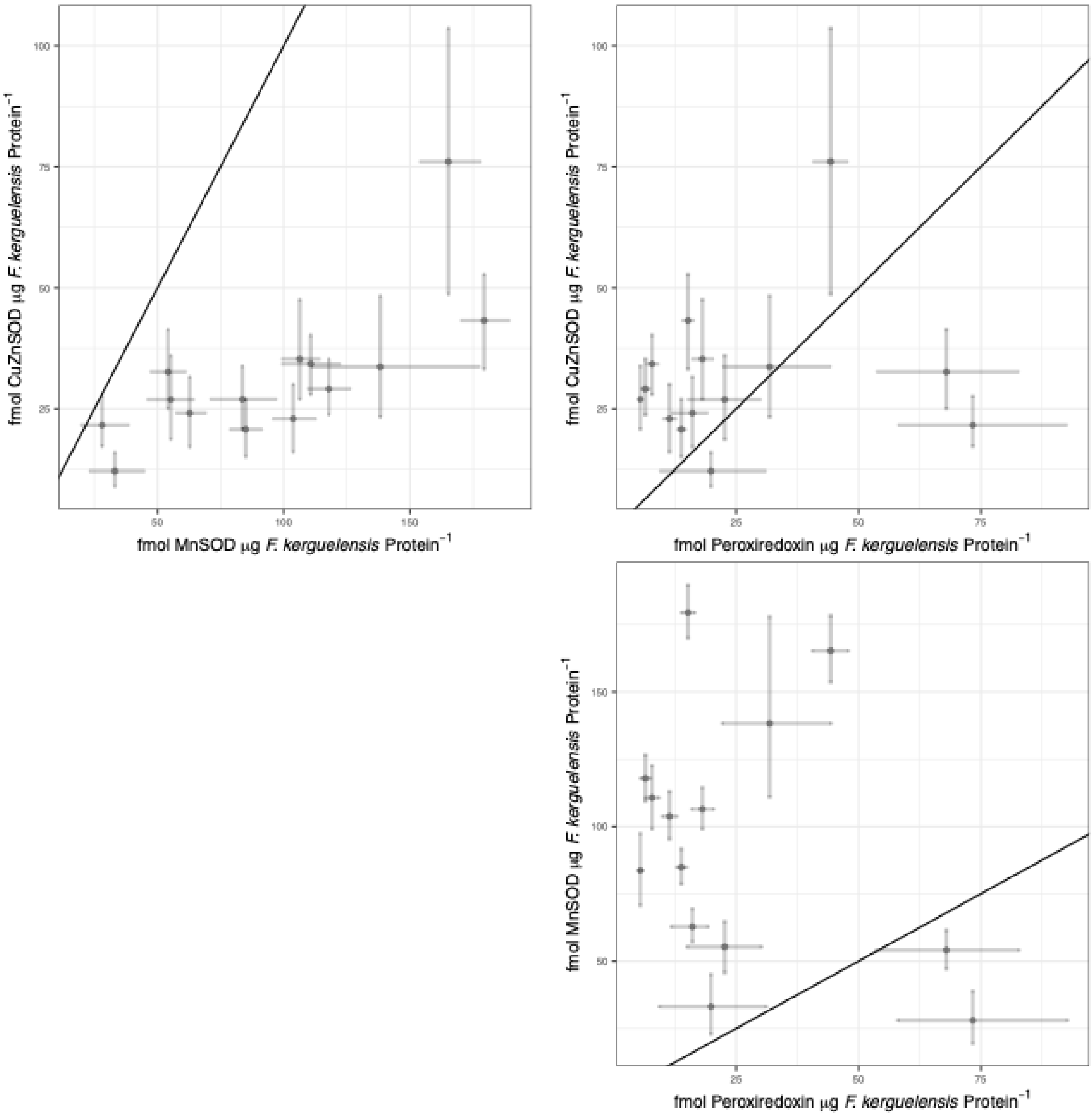


**Supplementary Figure S6**. Estimated *F. kerguelensis* protein abundances across the three proteins, MnSOD, CuZnSOD, and peroxiredoxin. Error bars refer to 95% credible intervals for both measurements. Black solid lines are the 1:1 lines.

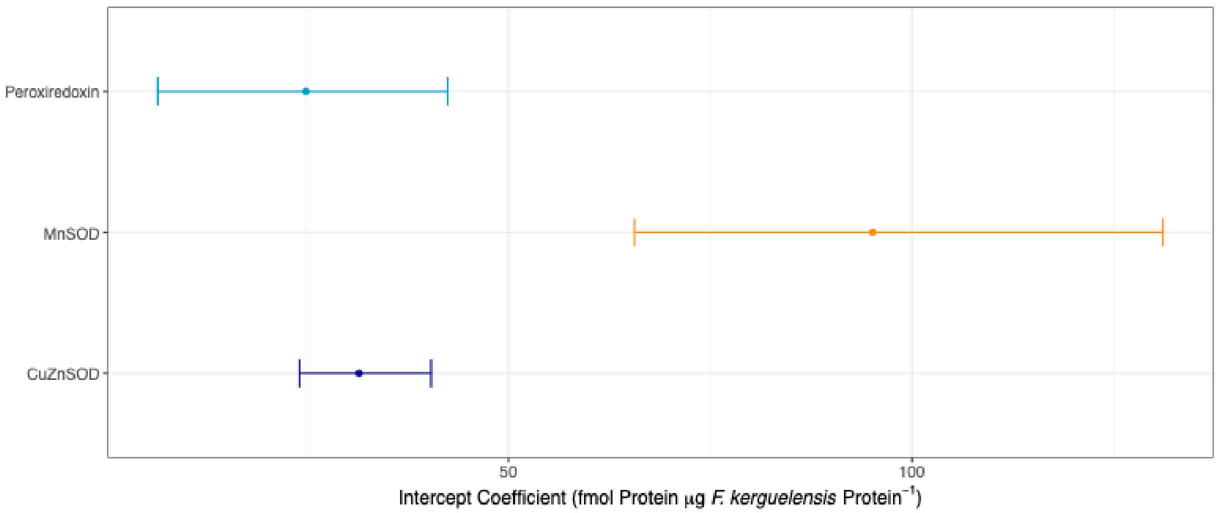


**Supplementary Figure S7.** Estimated intercept coefficients *F. kerguelensis* protein abundances.

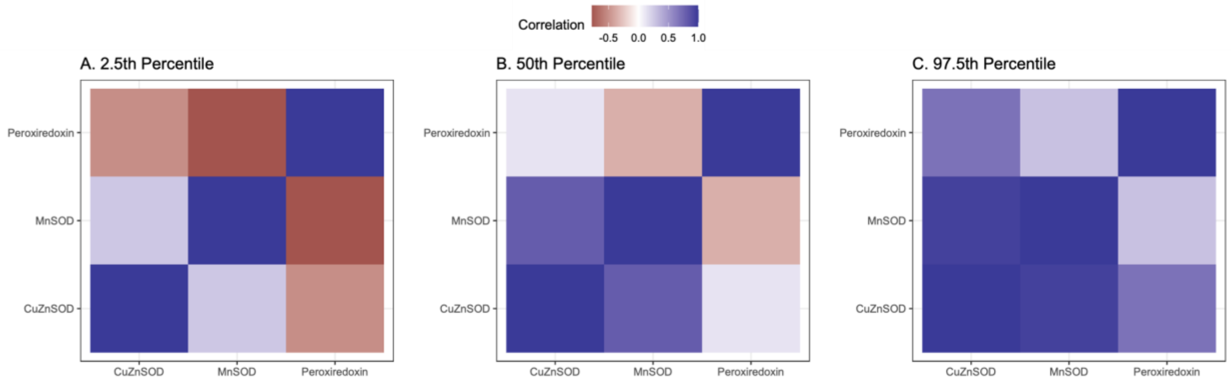


**Supplementary Figure S8.** Estimated correlation matrix that is used for modelling proteins as a multivariate normal distribution. (A) 2.5 percentile estimate, (B) median estimate, and (C) 97.5 percentile estimate.

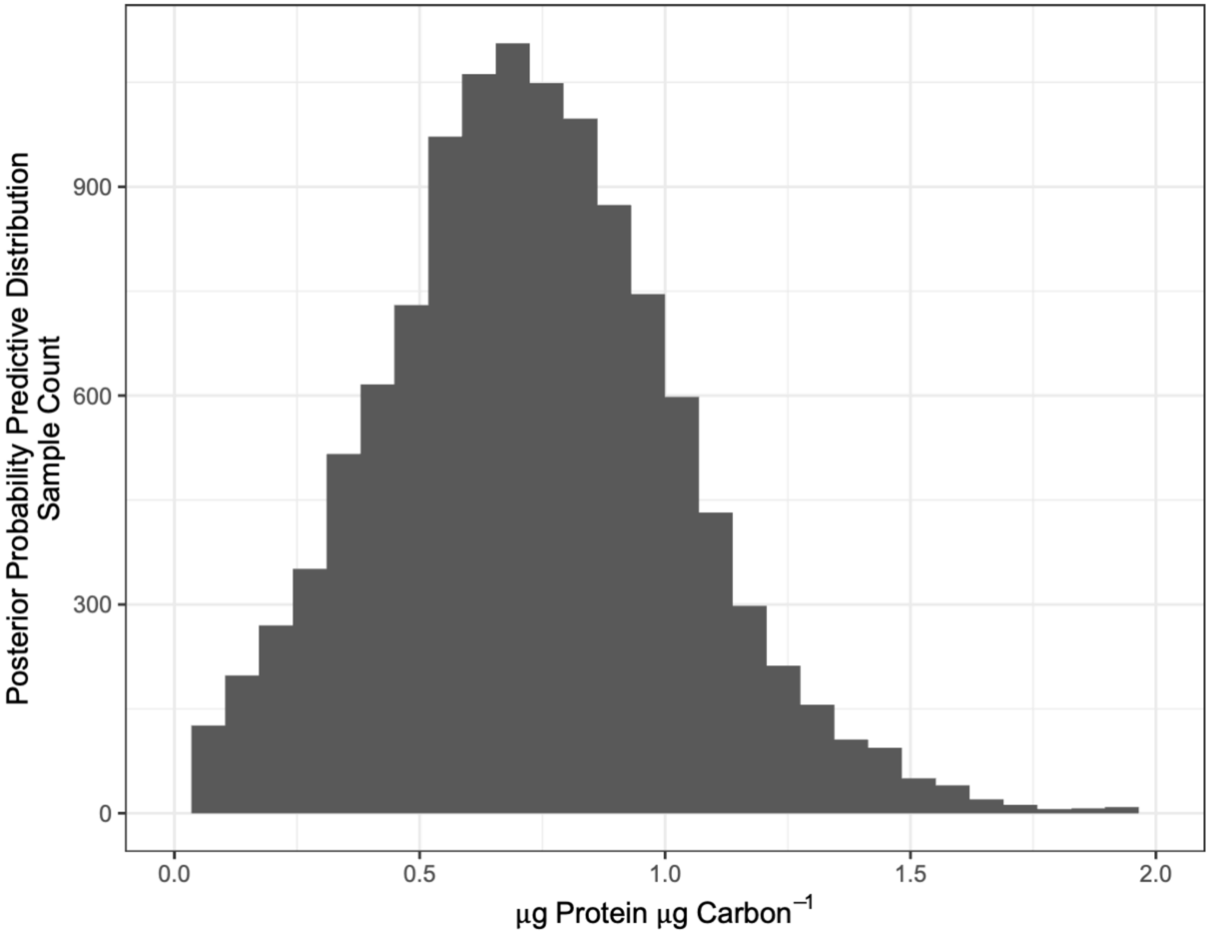


**Supplementary Figure S9**. Posterior predictive distribution for the protein-to-carbon mass ratio in *F. kerguelensis*.

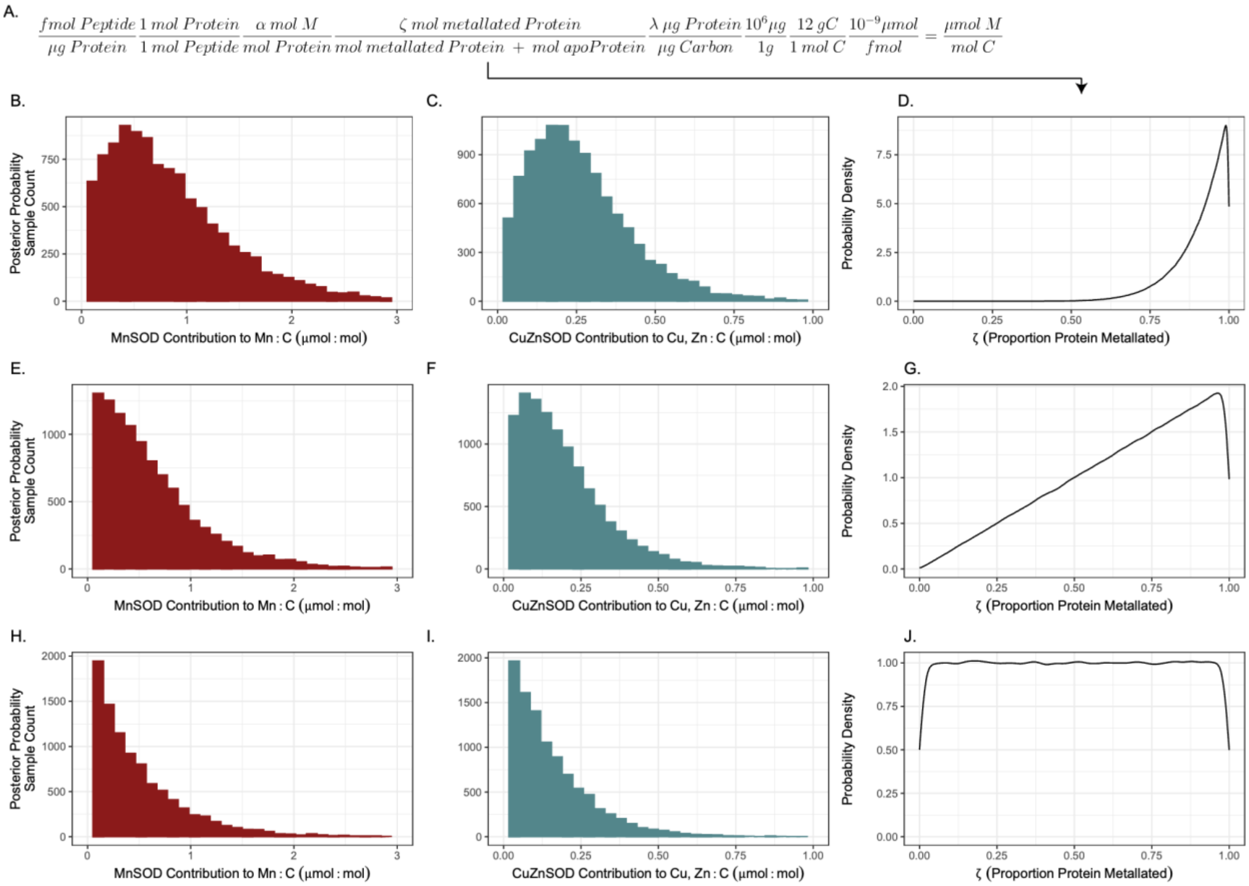


**Supplementary Figure S10**. Posterior predictions for contribution to metal:carbon assuming different metalation probability distributions (different rows). (A) Calculation for converting metaproteomic measurements into the contribution to metal (*M*) to carbon ratio (described in main text). (B, E, H) Posterior probability (the probability distribution after the data have been taken into account) samples showing the contribution of MnSOD to Mn:C ratio, assuming the protein is metalated by Mn. (C, F, I) Posterior probability samples showing the contribution of CuZnSOD to Cu, Zn:C ratios. (B-C) assume that the majority of protein present is metalated following the probability density plotted in (D), specifically a beta distribution with shape parameters equal to 10 and 1. (E-G) Posterior predictions for Mn:C assuming a beta probability distribution with shape parameters equal to 2 and 1 (G). (H-I) Posterior predictions for Mn:C assuming a beta probability distribution with shape parameters equal to 1 and 1 (J).

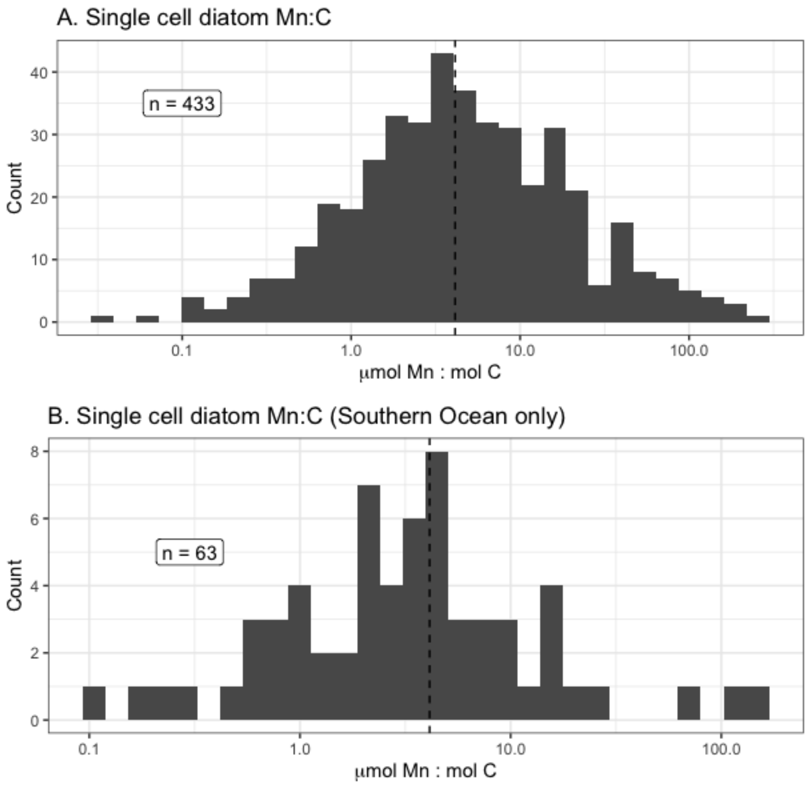


**Supplementary Figure S11**. Distribution of previously published values of Mn:C in single diatom cells from all available data (panel A) and data from the Southern Ocean (panel B).

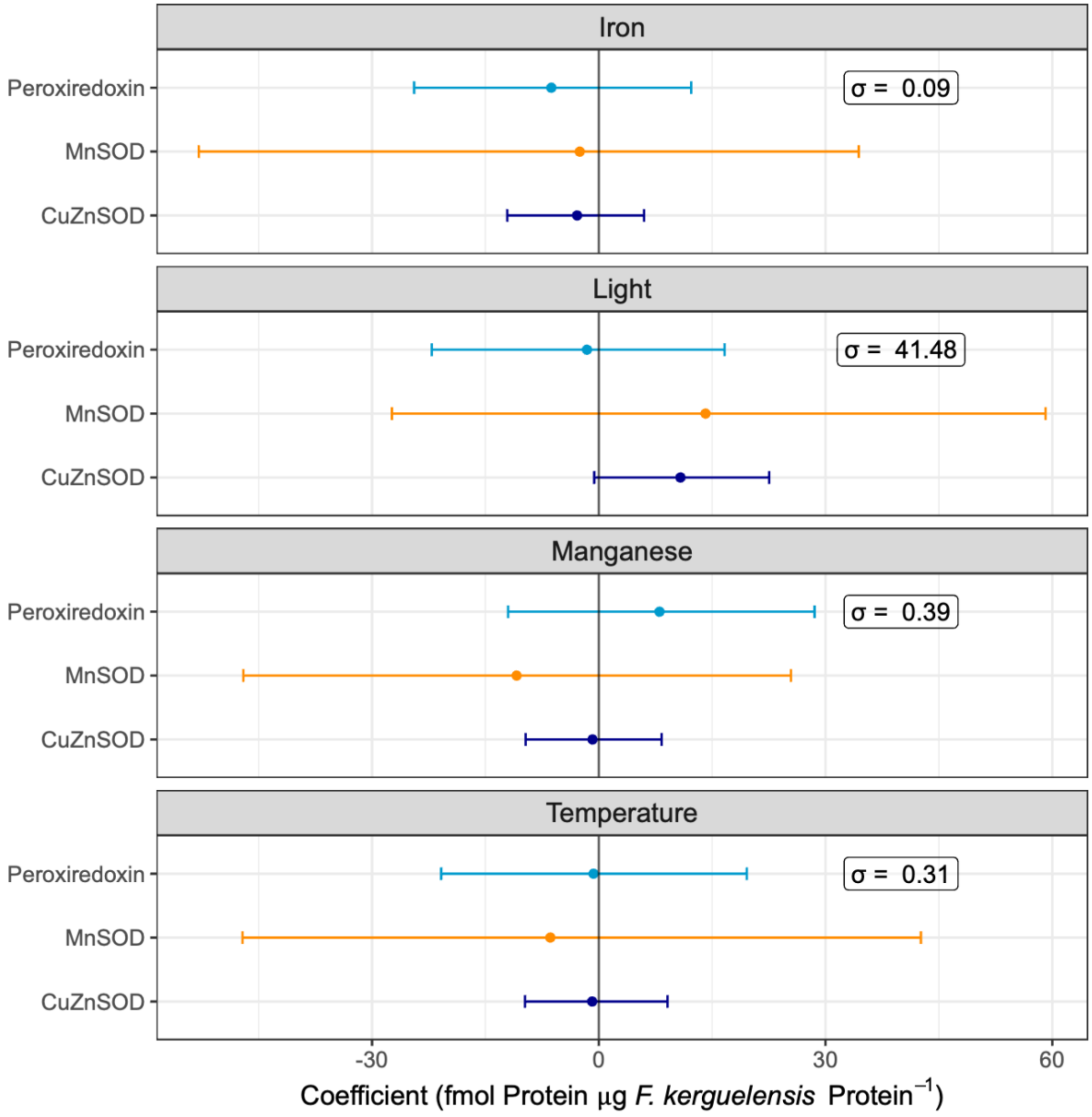


**Supplementary Figure S12**. Coefficient estimates (95% credible intervals shown) for four different environmental covariates across panels: iron, light, manganese, and temperature. Environmental covariates were z-score transformed prior to fitting the model, so the coefficients represent the association of one standard deviation change in the environmental covariate (observed standard deviations are shown in boxes) with change in individual proteins.

**Supplementary References**

1. Pino, L. K. *et al.* The Skyline ecosystem: Informatics for quantitative mass spectrometry proteomics. *Mass Spectrometry Reviews* **39**, 229–244 (2020).
